# Comparing LigandMPNN and Directed Evolution for Altering the Effector-Binding Site in the RamR Transcription Factor

**DOI:** 10.1101/2025.07.10.663684

**Authors:** Alia Clark-ElSayed, Ethan Creed, Kaila Nayvelt, Andrew Ellington

## Abstract

Recently, the number of ML-based tools for protein design has greatly expanded. Although there have been many successful uses of these tools for improved stability, solubility, and ligand binding, there have been fewer uses of these tools for designing proteins that have intrinsic allosteric mechanisms. In this regard, allosteric transcription factors (aTFs) are a class of regulatory proteins that includes repressors and activators that respond to environmental signals by allosteric communication to regulate their binding with DNA elements. The data exist for evaluating design algorithms for their ability to take allostery into account, as many aTFs have previously been engineered to respond to new ligands, enabling their use as biosensors. In particular, previous work from our lab used directed evolution to change the effector specificity of the transcriptional repressor, RamR, from cholic acids to each of five benzylisoquinoline alkaloids (BIAs). We wanted to see to what extent we could recapitulate these results by instead using LigandMPNN to design the ligand binding pocket. The wild-type RamR structure was predicted in complex with the five BIAs, and the binding pocket was then targeted for computational redesign. However, there was little overlap between the results of directed evolution and computational redesign, and in fact the nine redesigned protein variants tested proved not to be functional in *Escherichia coli*. Overall, these and other results suggest that different protein design methods may be needed to advance the computational design of allosteric or conformationally flexible proteins.

## Introduction

Bacteria have evolved to sense environmental and metabolic changes by detecting a variety of small molecule signals via allosteric transcription factors (aTFs)^1^. In general, the binding of small molecules changes the protein’s interactions with specific DNA sequences called operators, thereby controlling downstream biological processes^2^. Transcription factor-based biosensing has proven useful in a variety of biotechnology applications, including disease diagnostics^3^, contamination detection^4^, and monitoring of natural product biosynthesis^5^. Engineering transcription factors for targeted biosensing applications is of great importance, as it enables the development of specific and tunable detection systems.

RamR is a transcriptional repressor from the TetR family of regulator proteins from *Salmonella typhimurium*. RamR represses the expression of *ramR* and *ramA*, which increases the expression of the multidrug efflux system AcrAB-TolC^6^. RamR contains an N-terminal DNA binding domain and a C-terminal domain that interacts with ligands, altering the ability of the regulator to associate with DNA^7^. While RamR was initially discovered based on its role in affecting fluoroquinolone susceptibility in clinically isolated strains^8^, previous studies have also demonstrated the ability of RamR to respond to chemically diverse molecules, including berberine, crystal violet, dequalinium, ethidium bromide, Rhodamine 6G, and cholic acids^9,10^. This generality makes it an excellent test case for transcription factor engineering, and our lab has used a previously described method, Seamless Enrichment of Ligand Inducible Sensors (SELIS), to evolve RamR to respond sensitively and specifically to five different benzylisoquinoline alkaloids (BIAs)^11^.

The data in these studies serves as a remarkable test set for evaluating computational models, and in this work, we sought to test whether the deep learning-based model LigandMPNN might also be used to redesign the specificity of RamR. We first modeled RamR in a complex with the five ligands and used LigandMPNN to generate predicted aTFs for the fifteen residues originally randomized for directed evolution, primarily in the ligand-binding pocket. Little or no overlap between directed evolution and LigandMPNN designs was observed. To test the LigandMPNN designs, we cloned a number of variants into a cell-based reporter but found no sensors that were activated by their respective BIAs.

## Results

### Developing pipelines for computational prediction of transcription factors with altered effector specificities

Predicting functional complexes between small molecules and proteins is relevant to many protein engineering applications. Recently, deep-learning methods have advanced protein-ligand docking significantly^12,13^. However, a key challenge with docking is that it generally relies on a static picture of protein structure, whereas many structures undergo conformational changes upon binding. This is especially true for regulatory proteins, such as transcription factors. In addition, docking requires that a protein structure be available in the holo form: the conformation compatible with ligand binding. To take into account the structure of the docked, holo form of a protein, an important (indeed, saltatory) computational development is the all-atom version of RosettaFold (RFAA), which enables prediction of protein-ligand complexes, while still accounting for protein flexibility^14^. In initial RFAA demonstrations, when the structure of a protein was predicted with and without a ligand present, it proved able to predict known shifts in conformation, including domain movements, more subtle backbone movements, and flipping of side chain rotamers, suggesting that it could predict not only the docking of compounds, but potentially also the functional consequences of substitutions that might better enable docking^14^.

Other machine-learning methods have also emerged as powerful tools for engineering protein functionality, as well as structure^15^. ProteinMPNN is a deep learning-based method for inverse folding, which predicts a sequence that could fold into a desired structure^16^. Such inverse folding models have been demonstrated to generate highly stable (often too stable^17^) protein sequences^16^. More recently, LigandMPNN was released as an all-atom-aware version of ProteinMPNN^18^ that could be used to expand protein functionalities beyond just the 20 known amino acids, and researchers have in fact showed the ability to design sequences with nanomolar to micromolar affinity for steroidal, and folate compounds^18^.

We wanted to determine if some combination of RFAA and LigandMPNN could be used to modify the ligand-binding pocket of a transcription factor, including taking into account the necessary ‘gearing’ to achieve allosteric functionality. In previous work we used directed evolution to engineer RamR to respond sensitively and specifically to five benzylisoquinoline alkaloids (BIAs)^11^. Sensor engineering efforts centered on the compounds tetrahydropapaverine (THP), papaverine (PAP), rotundine (ROTU), glaucine (GLAU), and noscapine (NOS) (**Figure 1**). RamR showed very little responsivity to most of these BIAs, and libraries that spanned the five helices of the ligand-binding pocket were generated and selected for new ligand-binding functionality in *E. coli*. Ultimately, four of the five final sensor variants had EC_50_ values below for their respective ligands and also displayed >100-fold preference for their cognate BIAs.

**Figure 1.**
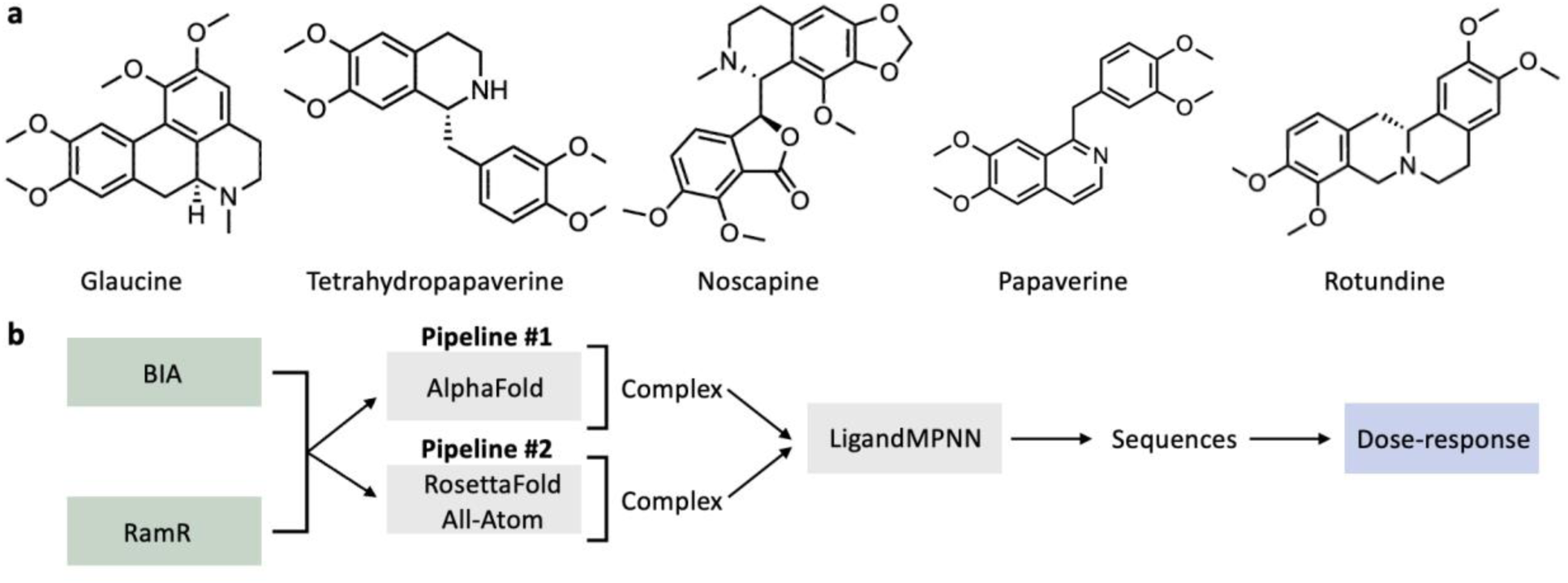
Using protein design tools to mutate the ligand binding pocket of RamR to bind BIAs. **(a)** Compounds used in this study: glaucine (GLAU), tetrahydropapaverine (THP), noscapine (NOS), papaverine (PAP), and rotundine (ROTU). **(b)** Computational design workflow. RamR-ligand complexes were predicted with AlphaFold and ligand docking and separately with RosettaFold All-Atom (RFAA). Structures were then designed with LigandMPNN, and sequences were cloned, expressed, and tested in *E. coli* for dose-response to the compound of interest.

Using these directed evolution methods as a ‘gold standard’ for comparison with computational design, we attempted to recapitulate them with RFAA and LigandMPNN via two conformational pipelines: 1) AlphaFold2 prediction of the structure of the transcription factor RamR, followed by ligand docking using DiffDock, then the use of LigandMPNN for altering effector binding; and 2) direct protein-ligand complex prediction for RamR using RFAA, following by LigandMPNN for altering effector binding (**Figure 1**). Between these pipelines, we hypothesized that RFAA might generate structures better suited for subsequent design with LigandMPNN, given its demonstrated accuracy in modeling protein-ligand complexes.

The original *Salmonella typhimurium* RamR protein was used as the basis for design, as was the case with directed evolution experiments. In Pipeline #1, this sequence was used to predict the protein structure using AlphaFold2^19^, and then DiffDock was employed to predict ligand binding, using SDF files as inputs. DiffDock generated multiple rank-ordered ligand conformations for the fixed protein structure, and for each BIA target we selected the highest-ranked ligand conformation and combined it with the protein structure into a single PDB file using PyMol (**Figure 2**). In Pipeline #2 the RamR sequence and SDF files of the five ligands were used as inputs for RFAA prediction of protein-ligand complexes (**Figure 3**).

**Figure 2.**
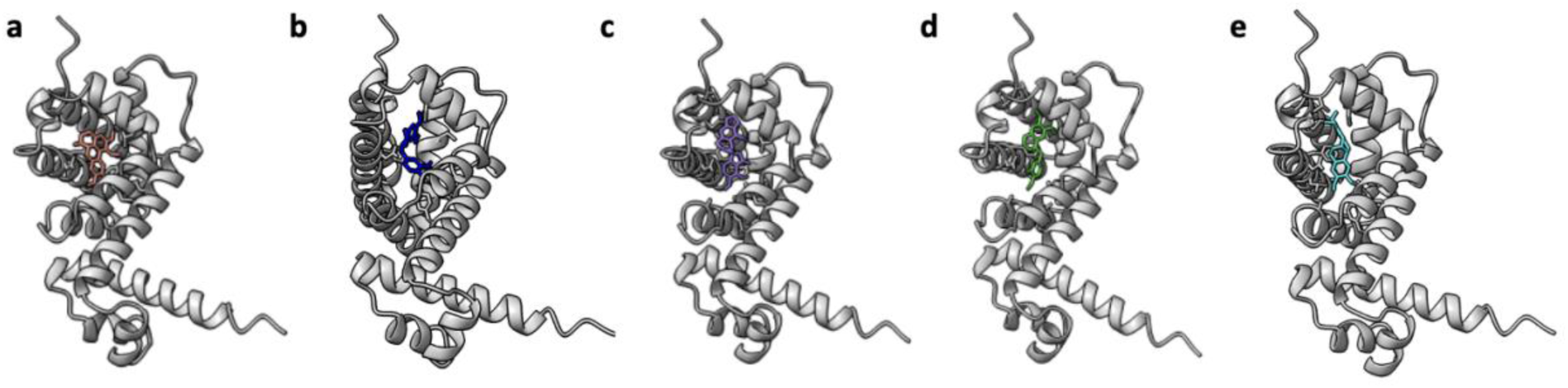
Predicted RamR-BIA complex structures generated through sequential AlphaFold2 protein structure prediction and DiffDock ligand binding prediction. **(a)** GLAU, **(b)** THP, **(c)** NOS, **(d)** PAP, **(e)** ROTU.

**Figure 3:**
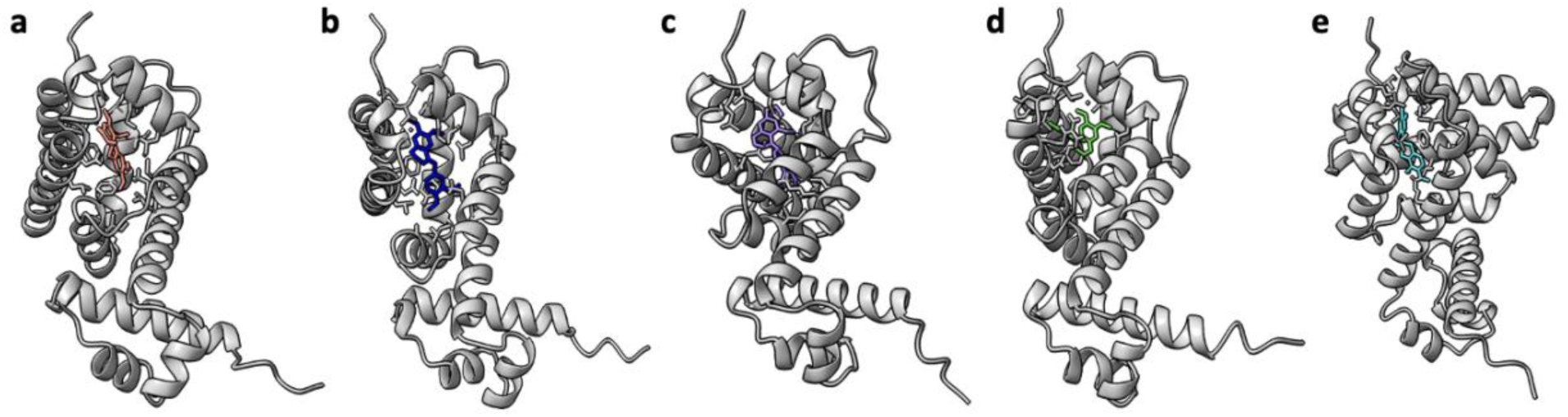
Predicted RamR-BIA complex structures generated with RFAA. (**a**) GLAU, **(b)** THP, **(c)** NOS, **(d)** PAP, **(e)** ROTU.

Significantly, all predicted ligand conformations from both methods localized to a binding pocket formed by five α-helices, which was in fact the previously characterized pocket that is known to accommodate various ligands, including ethidium bromide, rhodamine 6G, dequalinium, berberine, and crystal violet^20^. This is also fortuitous, given that the previously conducted directed evolution studies focused on randomizing these five helices.

### Using LigandMPNN to predict substitutions near the ligand binding pocket of RamR

To enable direct comparison between directed evolution and computational prediction, we used LigandMPNN to target the same 15 residues previously explored through site-saturation mutagenesis. LigandMPNN was run on 10 protein-ligand complexes: five of RamR with each of the five ligands modelled by AlphaFold and DiffDock (pipeline #1; 5 structures total) and five of RamR with each of the five ligands modelled by RFAA (pipeline #2; 5 structures total). Oddly, when pipeline #2 was used for predicting the RamR-ROTU complex, it output a structure that had the stereochemistry of the ligand swapped from R to S. We attempted multiple chemical structure files as input for RFAA; however, all the outputs had the stereochemistry swapped. In consequence, we excluded these designs from further consideration.

To our surprise, there was little overlap between the substitutions found by directed evolution and those predicted by LigandMPNN (**Figure 4**). Averaged across the 9 predicted sequences, the average number of mutations per protein was 9.0 (standard deviation: 1.4). This was superficially similar to the five known variants from directed evolution sequences, where the average number of mutations per sequence was 9.4 (standard deviation: 2.3). However, LigandMPNN failed to recapitulate the results of the directed evolution campaign in many regards. For all five BIA ligands, there were multiple positions where the directed evolution variant contained a substitution, but the LigandMPNN designs did not, and vice versa. Through the entire set of predictions, there is not a single residue where directed evolution and both LigandMPNN designs agreed on the identity of a substitution.

**Figure 4.**
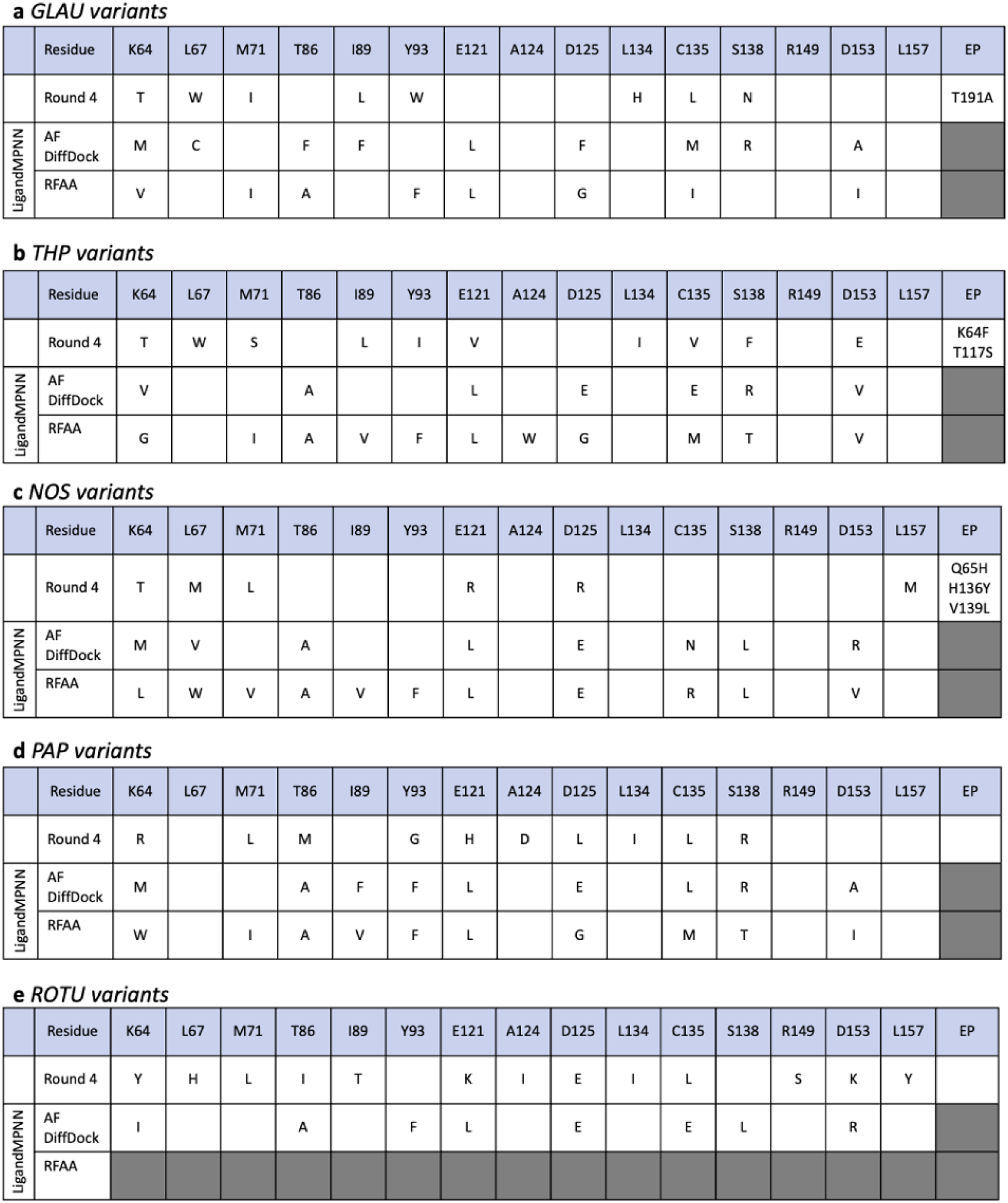
Variants for comparing directed evolution and LigandMPNN methods for transcription factor engineering. **(a)** GLAU, **(b)** THP, **(c)** NOS, **(d)** PAP, **(e)** ROTU. Row 1 of each chart is the variants from the round 4 directed evolution. Rows 2 and 3 are the LigandMPNN-designed sequences for each protein-ligand complex prediction method.

More liberally, there are four other positions in which at least one of the machine learning methods identified the same substitution as directed evolution, and there were 11 instances in which both directed evolution and machine learning declined to mutate the original wild-type protein. Interestingly, there are specific amino acids that were frequently used by LigandMPNN (glutamic acid, alanine, valine, and phenylalanine, **Supplementary Table 1**), relative to directed evolution. This likely reflects bias in the underlying training sets for the algorithm and further emphasizes the fact that actual evolutionary trajectories are likely more robust than machine learning approaches.

### LigandMPNN fails to generate protein variants that respond to BIAs

Of course, it is also possible that machine learning methods have yielded functional transcription factors for new effectors but took different paths through sequence space to do so than evolution. To test this hypothesis, the LigandMPNN-designed sequences were cloned into a reporter vector in *Escherichia coli* (**Supplementary table 1**). This reporter vector not only expresses a given RamR variant but also contains GFP under the control of a RamR operator and is similar to reporter vectors we have used in previous biosensor studies^21^ (**Figure 5**). Cells expressing the designed RamR variants were induced with increasing concentrations of each BIA, and the resultant fluorescence measured. In all cases, the transcription factors derived by directed evolution were functional, showing induction, while the sequences designed with LigandMPNN showed no induction (**Figure 6**). Interestingly, all designed variants except for one had high background signal in the absence of the ligand, suggesting that the protein is no longer able to tightly bind the operator sequence.

**Figure 5.**
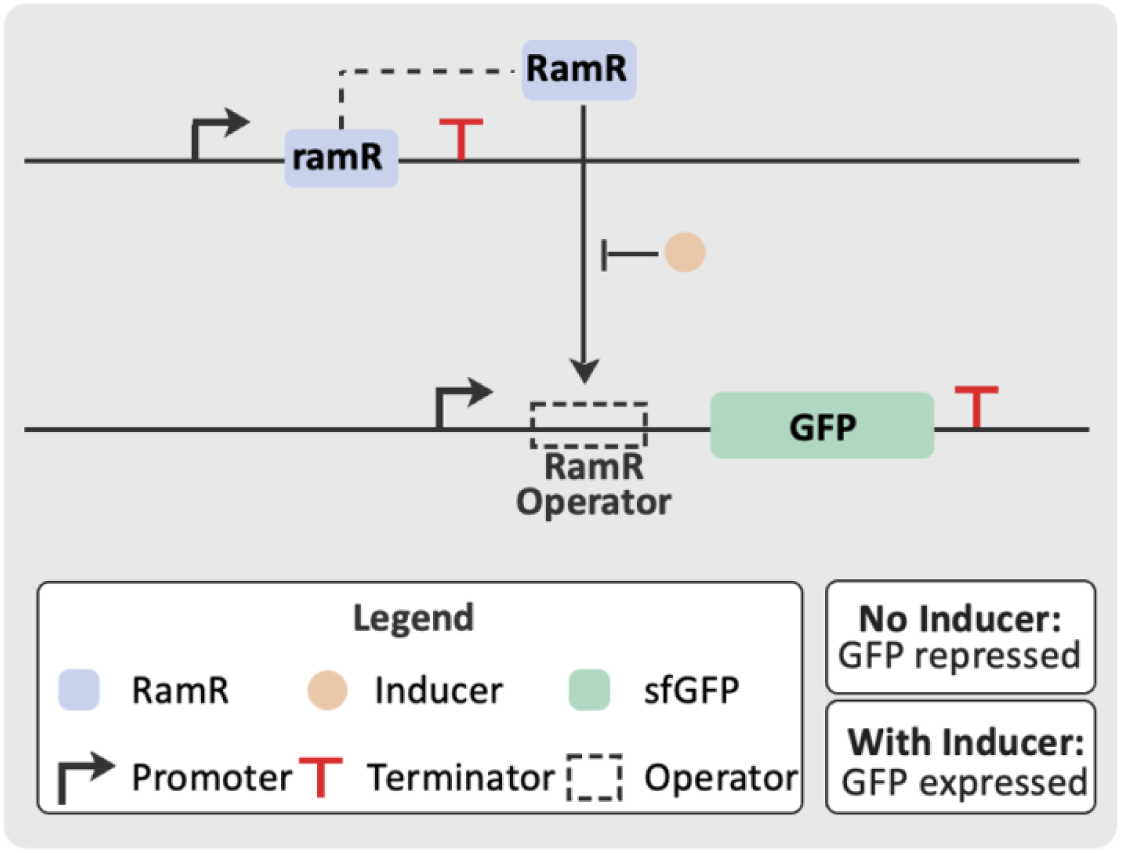
Genetic construct for dose-response measurements. The top panel describes the assay condition where the ligand has not been added to the cells. RamR binds to the specific operator sequence upstream of GFP, preventing the transcription of the reporter gene by RNA polymerase. The bottom panel describes the assay condition where the compound has been added to the cells. RamR binds to the compound, resulting in a conformation change which causes RamR to release from the operator sequence. RNA polymerase is permitted to transcribe the gene encoding GFP, resulting in an increase in fluorescence.

**Figure 6.**
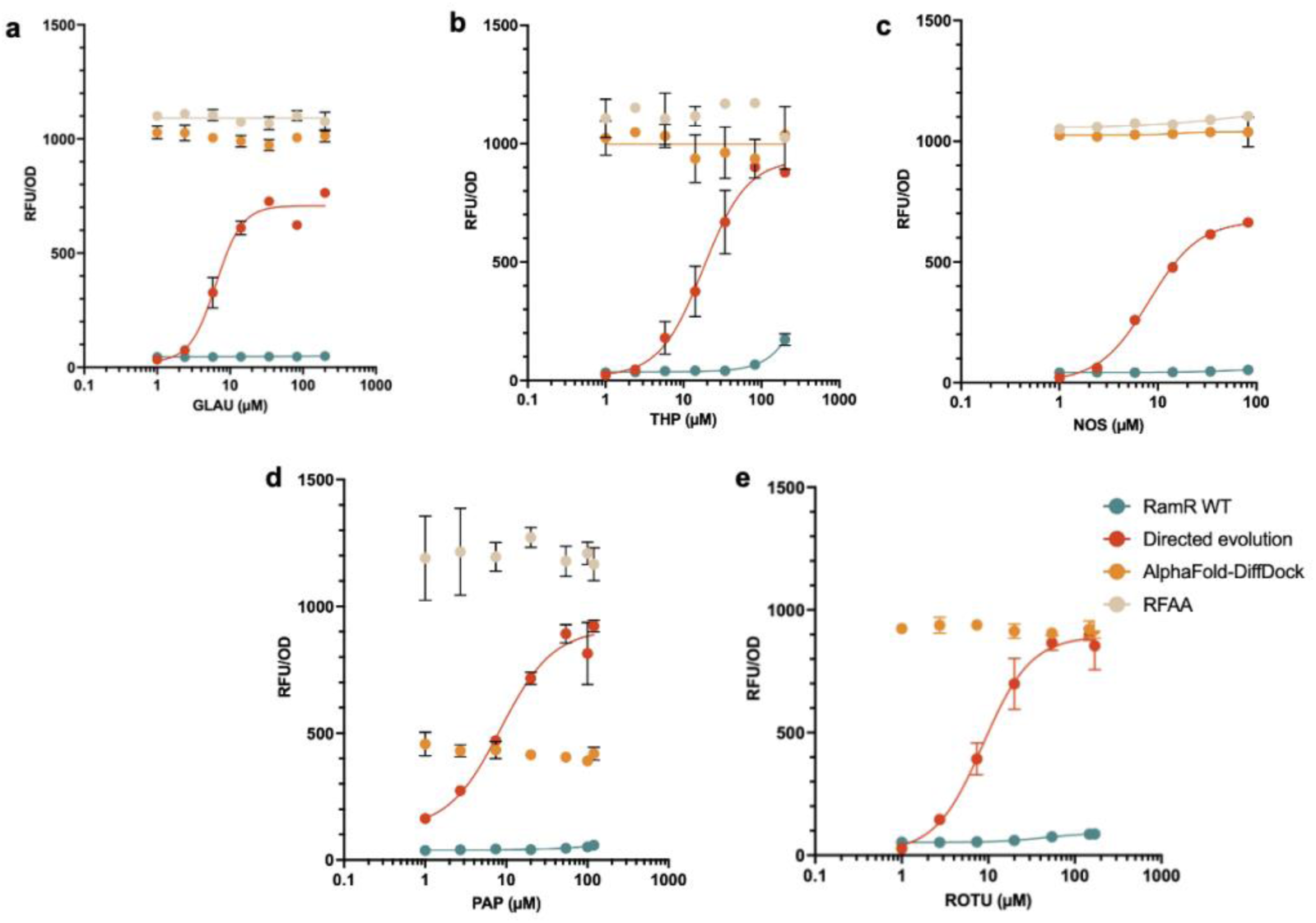
Dose-response measurements from induction assay. **(a)** GLAU, **(b)** THP, **(c)** NOS, **(d)** PAP, **(e)** ROTU. Color legend for variants: green; wild type, red; directed evolution, orange; LigandMPNN variants predicted from AlphaFold-DiffDock structure, beige; LigandMPNN variants predicted from RFAA structure.

## Discussion

Herein, we compare two methods for altering the effector-binding domain of the repressor RamR: previous directed evolution methods that successfully altered the binding pocket while retaining transcription factor function, and machine learning methods based on docking via either DiffDock or RFAA, and subsequent mutation by LigandMPNN. Should these computational models work for the successful prediction of new biosensors it would be a great advance, since crystal structures of protein targets, especially with small molecule ligands, are often unavailable. However, not only did the modeled variants not recapitulate the successes of directed evolution, but they also showed no induction in the presence of ligands.

Indeed, the predicted substitutions seem to ‘break’ the allostery of the transcription factor in general by enforcing a conformation that no longer binds to DNA. Similarly, another recent evaluation of ProteinMPNN found that designed variants of NanoLuc had improved stability and solubility, but lost appreciable enzymatic activity^17^. There are many studies that suggest that stability and conformational specificity limit the flexibility necessary for enzyme function^22,23,24^. By apparently focusing on designs that primarily stabilize structures, LigandMPNN fails to predict or loses interactions necessary for allostery and / or catalysis. Overall, while these results further demonstrate limitations in computational design tools, they also suggest that comparisons with directed evolution experiments can be used as a gold standard for assessing (and ultimately improving) such tools. In this regard, the data sets describing the directed evolution of RamR for the various benzylisoquinoline alkaloids may prove extremely valuable for training evolution-aware fitness models.

## Methods

### Structure prediction

For previous directed evolution work, the protein sequence for RamR was retrieved from the NCBI database (PDB: 3VVX_A)^25^.

For the AlphaFold model, the RamR sequence was folded using ColabFold v1.5.5: AlphaFold2 using MMseqs2(pLDDT=92.8, pTM=0.868)^19^. For four of the five BIAs, SDF files of 3D conformer structures were retrieved from the Pubchem database (GLAU: 16754, NOS: 275196, PAP: 4680, THP: 5418)^26^. For ROTU, there did not exist a 3D conformer in the Pubchem database, therefore the smile string (COC1=C(OC)C2=C(C=C1)C[C@]3([H])C4=CC(OC)=C(OC)C=C4CCN3C2) was converted to a 3D structure using NovoPro Lab’s online tool^27^. The AlphaFold model was then used for ligand docking with DiffDock^28^. The output SDF files from DiffDock were combined with the AlphaFold PDB structure using PyMol, and each of the protein-ligand complexes was exported as PDB files.

For the second complex prediction, the RamR sequence was predicted in a small molecule complex with the same SDF files described previously. RFAA was run using the base config as the default^29^.

### LigandMPNN

All of the protein-ligand complexes were predicted using LigandMPNN^18^. The run parameters were as follows: model type= ligand_mpnn, seed= 111, Redesigned residues= “A64 A67 A71 A86 A89 A93 A121 A124 A125 A134 A135 A138 A149 A153 A157”, Ligand_mpnn_use_side_chain_context=1.

### Strains, plasmids, and media

*E. coli* DH10B was used for the characterization of all biosensors. The plasmids described in this work were ordered as clonal genes (plasmids) from Twist Biosciences. The plasmid sequence used for assays is provided in Supplementary Table 1. LB with 1.5% agar and kanamycin 50 (µg/mL) was used for selection after the transformation of the plasmids described in this work. LB Miller (LB) medium was used to grow all cells. When kanamycin was added to the culture medium, its final concentration was 50µg/mL. Super Optimal broth with Catabolite repression (SOC) medium was used in plasmid transformations to recover cells before plating.

### Benzylisoquinoline alkaloids

Cells were induced with the following chemicals: THP (TCI Chemicals, N0918), PAP (MP Biomedicals, 19026105), GLAU (Carbosynth, ACA72232), ROTU (Alfa Aesar, AAJ6332803), NOS (Sigma-Aldrich, 363960).

### Dose-response measurements

10µl of chemically competent DH10B cells (New England Biolabs) were transformed with 1µl of plasmid, resuspended in 100µl of SOC, and then recovered at 37°C for 1 hour before plating on LB/Agar with kanamycin. Following growth at 37°C overnight, triplicate colonies were inoculated into 900µl LB containing kanamycin in a 96-deep-well round bottom plate (ThermoScientific, 278752) sealed with an AeraSeal film (Excel Scientific, BS-25). Plates were incubated at 37°C and 1000 r.p.m in a shaking incubator (VWR, Incubating Mini Shaker) for 16 hours.

20µl of the overnight growth was inoculated into 900µl of LB containing kanamycin in a 96-deep-well plate, sealed with an AeraSeal film, and grown at 37°C with shaking (1000 r.p.m) for two hours. Cultures were then induced with the compound of interested dissolved DMSO (Sigma-Aldrich, D8418), DMSO, and LB medium. The final volume was 1000µl per well, with a final DMSO concentration of 1%, and variable concentration of the target ligand. Cells were grown at 37°C with shaking (1000 r.p.m) for four hours, after which cells were pelleted in a centrifuge (Eppendorf, 5810R) at 3000xg for 10 minutes. The supernatant was removed, and cell pellets were resuspended in 900µl of phosphate-buffered saline (PBS, prepared from Gibco PBS (10X), pH 7.4, 70011069).

100µl of the resuspension was transferred to a 96-well black microplate with a clear flat bottom (Corning, 3631), from which the fluorescence (excitation, 475nm; emission, 509nm; gain, 50) and absorbance (600nm) were measured using the Tecan infinite 200 Pro plate reader.

## Statistical analysis and reproducibility

All data in the text are displayed as mean ± standard deviation from three biological replicates. The data was fitted using the [Agonist] vs. response (four parameters) fit of the GraphPad Prism 10.0 software^30^.

## Author contributions

A.C.E designed the experiments, performed the analysis, and wrote the manuscript. E.C. and K.N. performed the dose-response characterization assays. A.E. supervised the study.

## Acknowledgements

Research reported in this publication was supported by the National Institute Of General Medical Sciences of the National Institutes of Health under Award Number T32GM139796 to ACE and 1R01 GM146093-01 A1 to A.D.E. The content is solely the responsibility of the authors and does not necessarily represent the official views of the National Institutes of Health.

This study was funded by the Defense Advanced Research Projects Agency (DARPA) under award number N660012324041. The views, opinions, and/or findings expressed are those of the authors and should not be interpreted as representing the official views or policies of the Department of Defense or the U.S. Government.

The authors acknowledge financial support from the Welch Foundation (F-1654).

The author wishes to thank Professor Edward Marcotte for his guidance in conceptualizing the computational framework of this research.

## Supplementary information

**Supplementary table 1.**
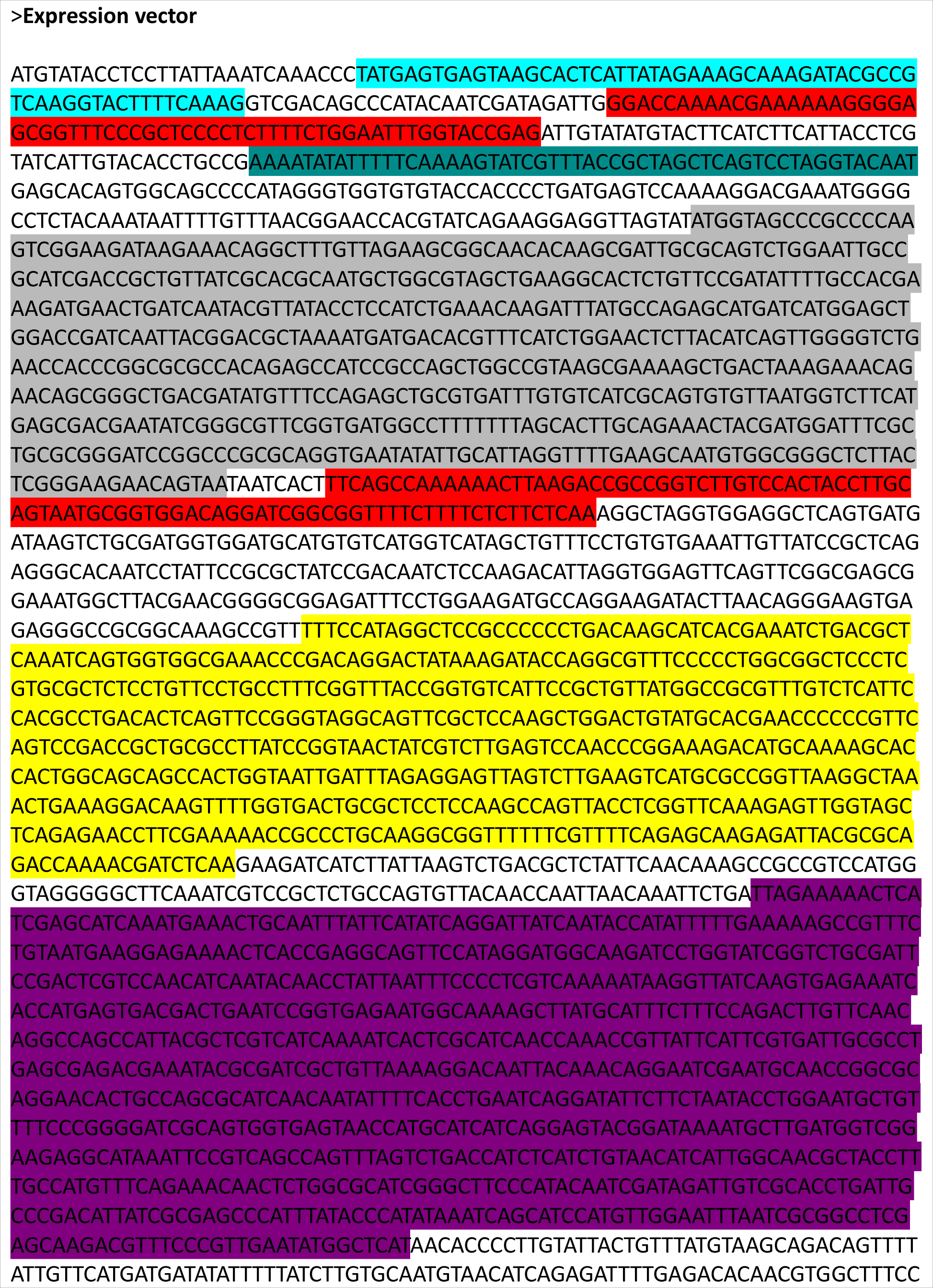
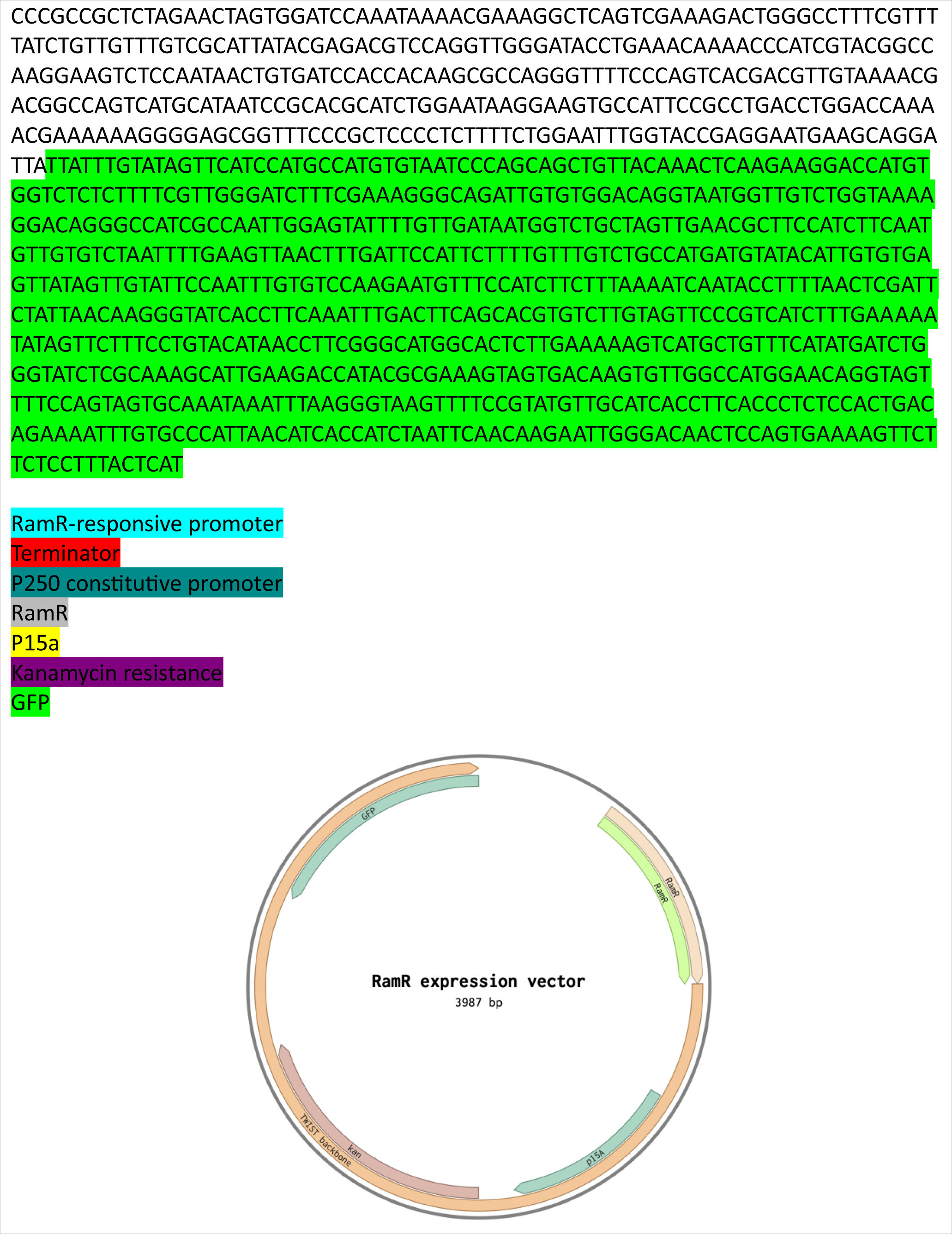
Expression plasmid sequence. Components of the plasmid are highlighted, and a map of the plasmid is provided at the bottom of the table.

**Supplementary table 2.**
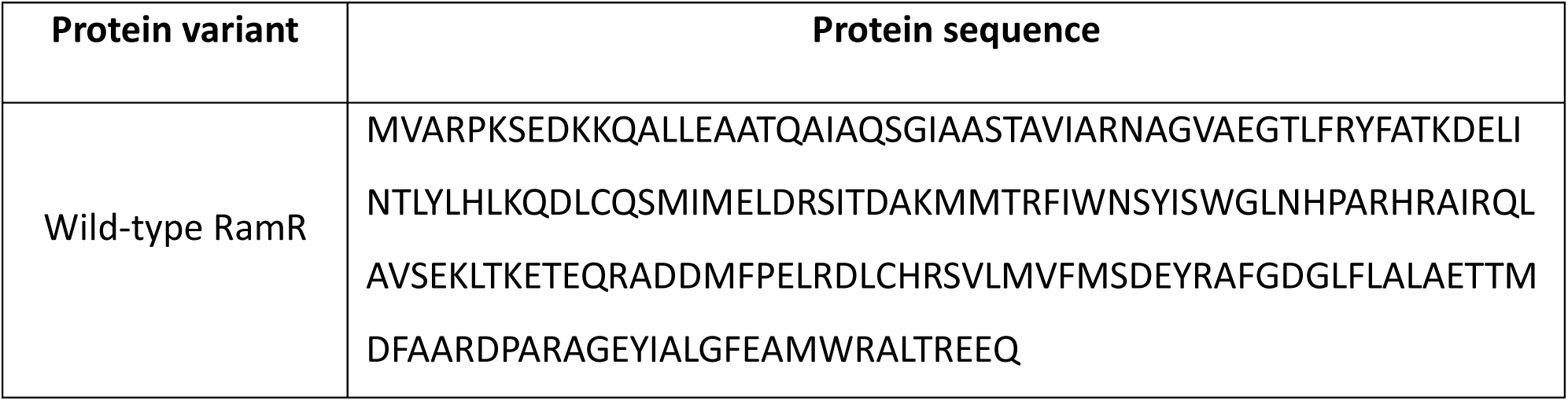
Wild-type RamR protein sequence.

**Supplementary table 3.**
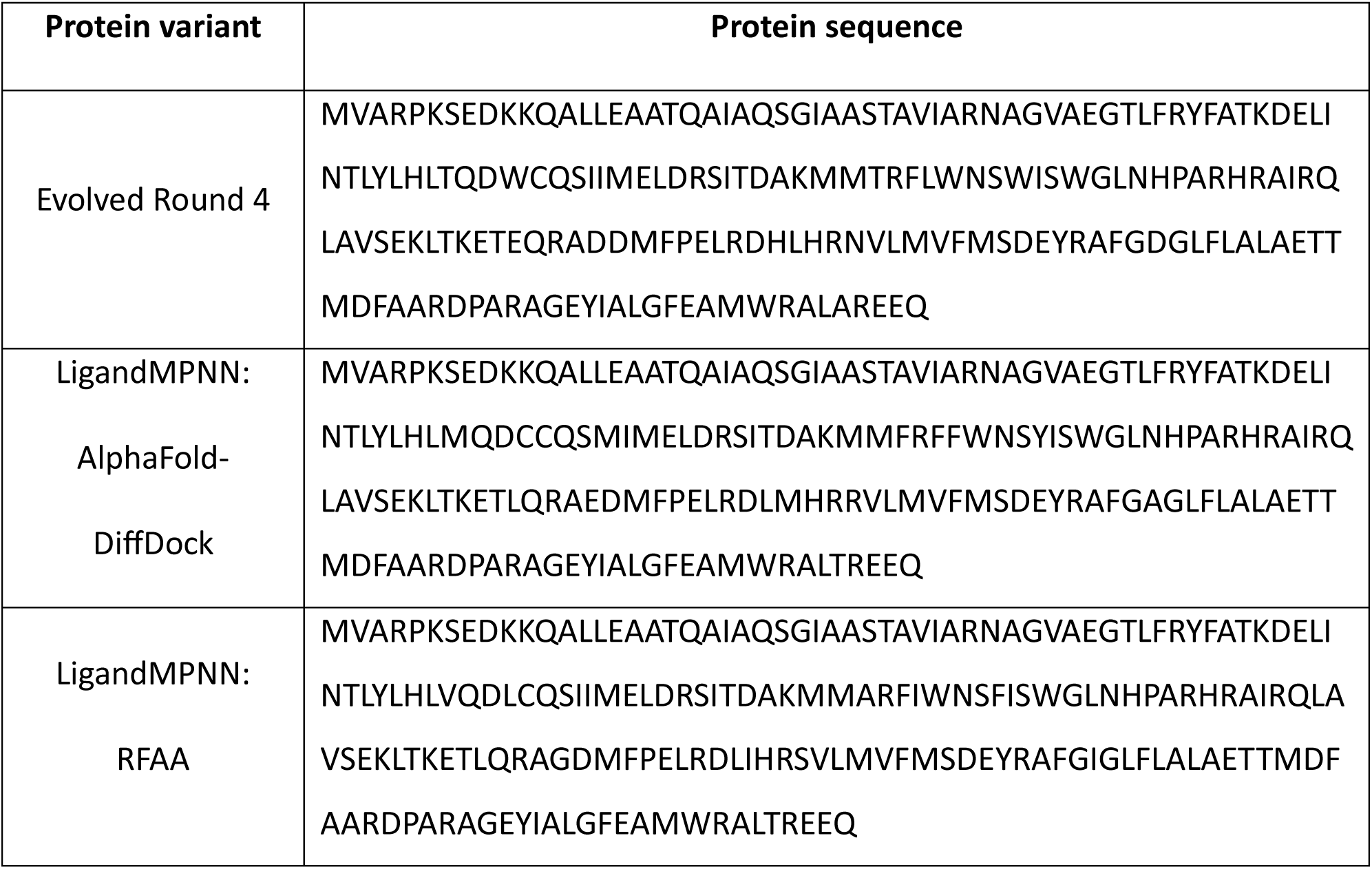
GLAU protein variants.

**Supplementary table 4.** THP protein variants.

| Protein variant | Protein sequence |
| --- | --- |
| Evolved Round 4 | MVARPKSEDKKQALLEAATQAIAQSGTAASTAVIARNAGVAEGTLFRYFATKDELI<br>NTLYLHLFQDWCQSSIMELDRSITDAKMMTRFLWNSIISWGLNHPARHRAIRQL |
|  | AVSEKLSKETVQRADDMFPELRDIVHREVLMVFMSDEYRAFGEGLFLALAETTM<br>DFAARDPARAGEYIALGFEAMWRALTREEQ |
| LigandMPNN:<br>AlphaFold-DiffDock | MVARPKSEDKKQALLEAATQAIAQSGIAASTAVIARNAGVAEGTLFRYFATKDELI<br>NTLYLHLVQDLCQSMIMELDRSITDAKMMARFIWNSYISWGLNHPARHRAIRQL<br>AVSEKLTKETLQRAEDMFPELRDLEHRRVLMVFMSDEYRAFGVGLFLALAETTM<br>DFAARDPARAGEYIALGFEAMWRALTREEQ |
| LigandMPNN:<br>RFAA | MVARPKSEDKKQALLEAATQAIAQSGIAASTAVIARNAGVAEGTLFRYFATKDELI<br>NTLYLHLGQDLCQSIIMELDRSITDAKMMARFVWNSFISWGLNHPARHRAIRQL<br>AVSEKLTKETLQRWGDMFPELRDLMHRTVLMVFMSDEYRAFGVGLFLALAETT<br>MDFAARDPARAGEYIALGFEAMWRALTREEQ |

**Supplementary table 5.**
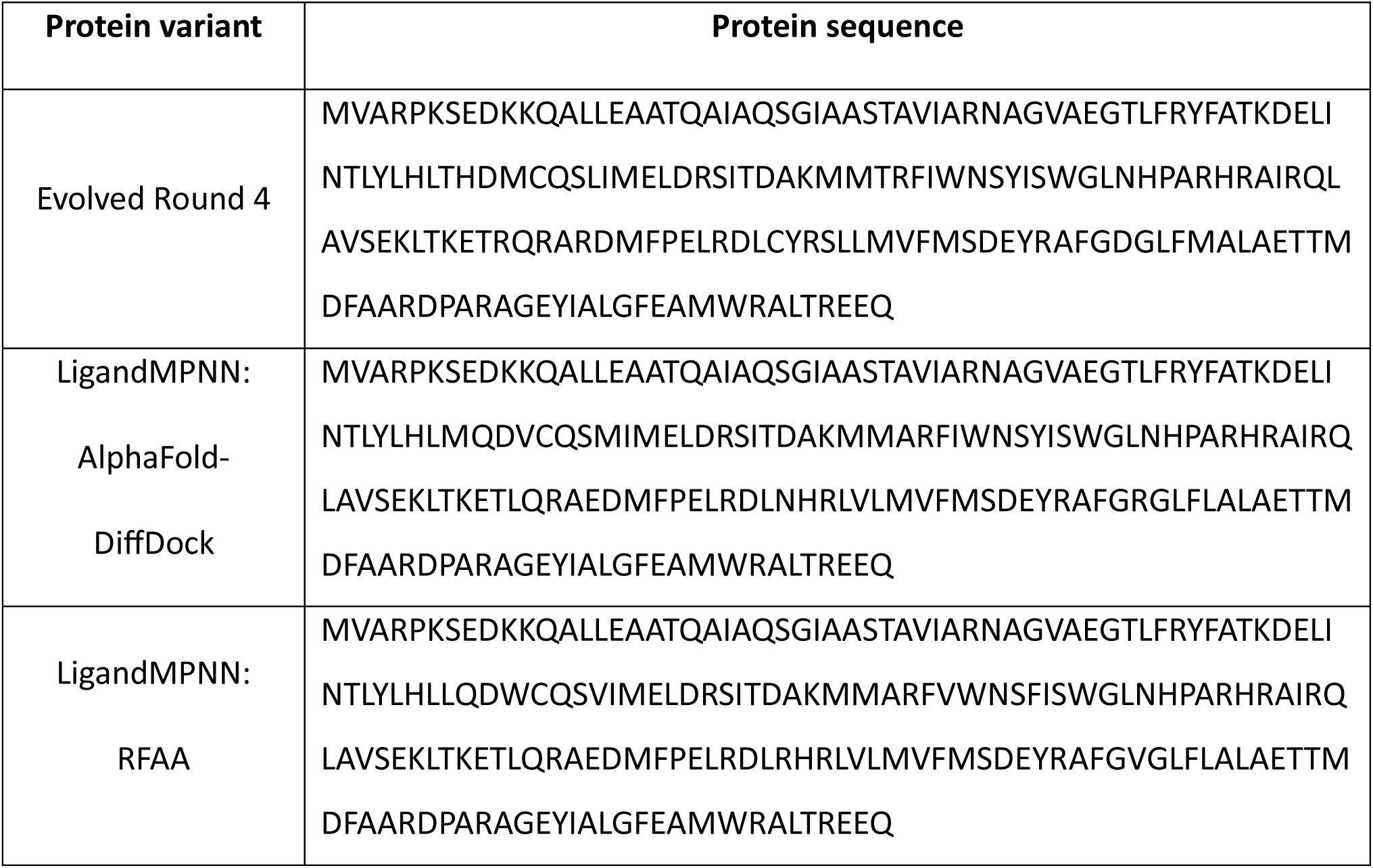
NOS protein variants.

**Supplementary table 6.**
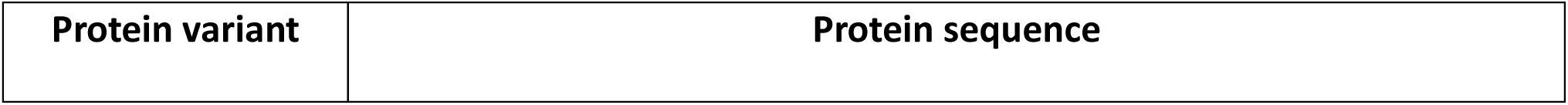
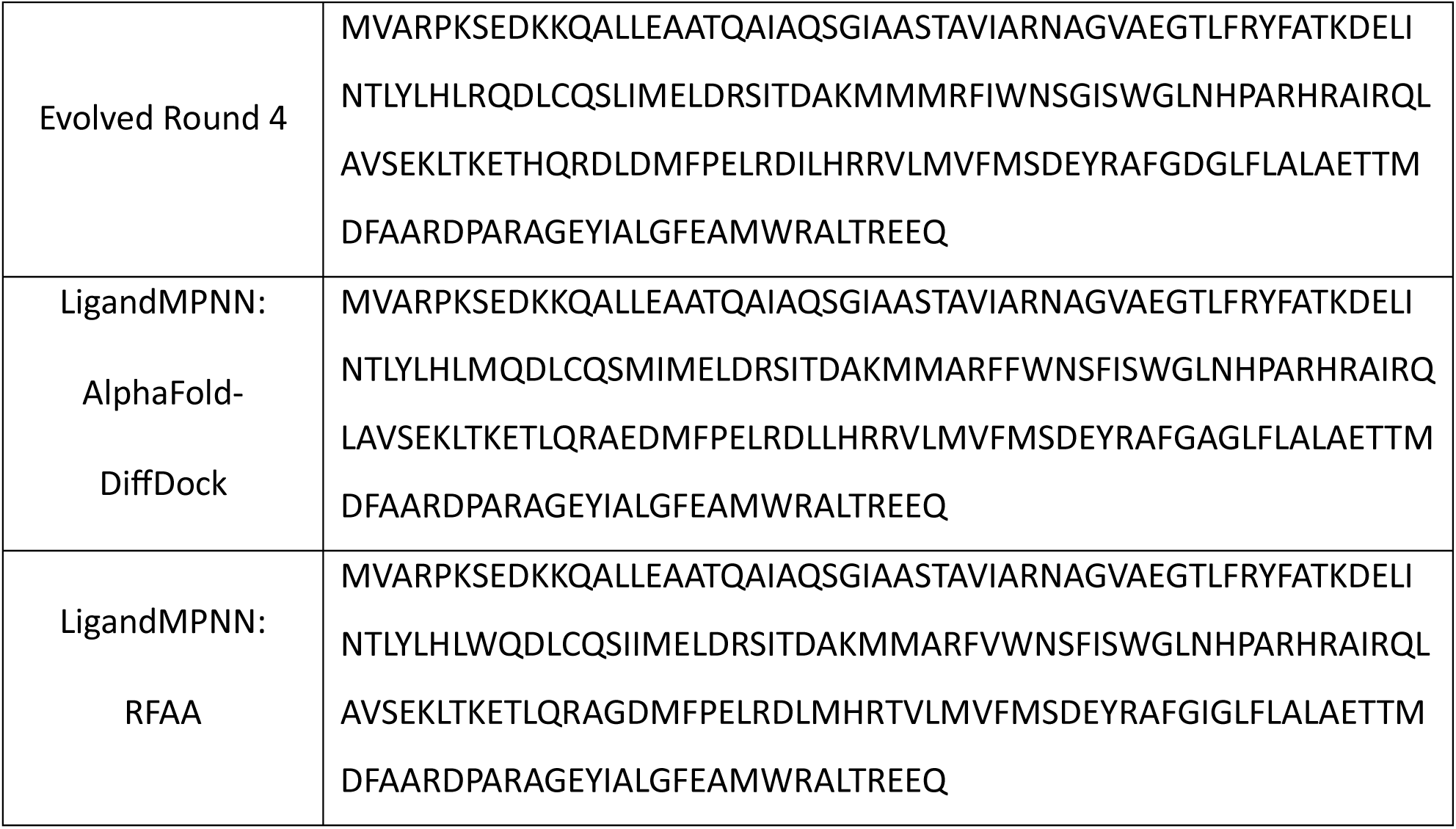
PAP protein variants.

**Supplementary table 7.**
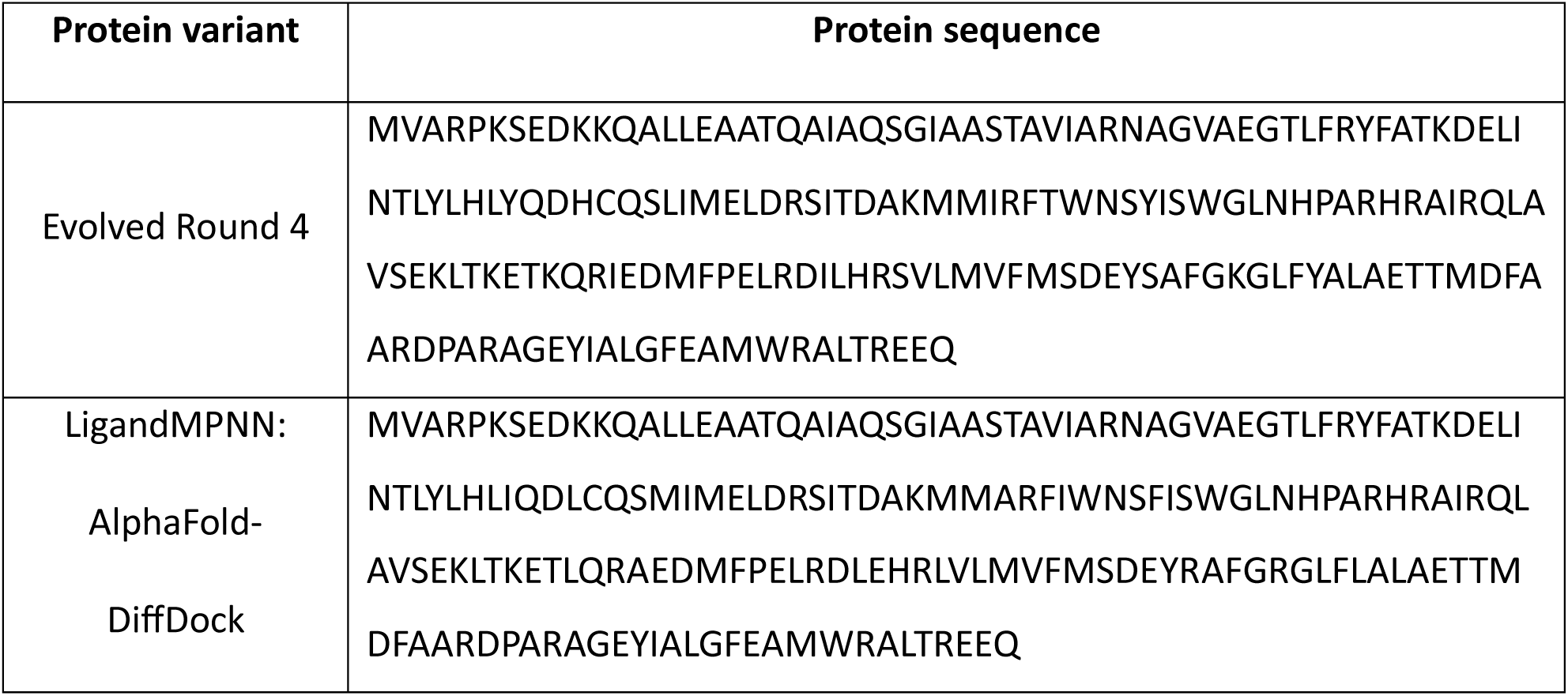
ROTU protein variants.

**Supplementary table 8.** Amino acid distribution of designed residues.

| Amino Acid | Round 4 Directed Evolution<br>variants | LigandMPNN design variants |
| --- | --- | --- |
| R | 4 | 6 |
| H | 3 | 0 |
| K | 2 | 0 |
| D | 1 | 0 |
| E | 2 | 7 |
| S | 2 | 0 |
| T | 4 | 2 |
| N | 1 | 1 |
| Q | 0 | 0 |
| C | 0 | 1 |
| G | 1 | 4 |
| P | 0 | 0 |
| A | 0 | 10 |
| V | 2 | 10 |
| I | 7 | 7 |
| L | 9 | 14 |
| M | 3 | 6 |
| F | 1 | 10 |
| Y | 2 | 0 |
| W | 3 | 3 |

